# Direct electrical stimulation of the human amygdala enhances recognition memory for objects but not scenes

**DOI:** 10.1101/2025.06.05.658078

**Authors:** Krista L. Wahlstrom, Justin M. Campbell, Martina K. Hollearn, James Swift, Markus Adamek, Lou Blanpain, Tao Xie, Tyler Davis, Peter Brunner, Stephan B. Hamann, Amir Arain, Lawrence N. Eisenman, John D. Rolston, Shervin Rahimpour, Joseph R. Manns, Jon T. Willie, Cory S. Inman

## Abstract

Research from neuroscience studies using invasive neuroanatomy-inspired direct manipulations suggests the basolateral amygdala (BLA) mediates a generalized modulation of many different types of memory. In contrast, noninvasive, psychology-inspired indirect correlations suggest that specificity exists in how the BLA prioritizes experiences in memory. We used direct electrical stimulation of the BLA to investigate the specificity of the memory enhancement in the human brain. Patients undergoing intracranial monitoring via depth electrodes viewed object and scene images, half of which were followed by BLA stimulation. Stimulation enhanced long-term memory for object but not scene images. Furthermore, BLA stimulation elicited stronger evoked responses in the anterior vs. posterior medial temporal lobe (MTL), regions that preferentially process object and scene learning, respectively. These results suggest the BLA exerts an important influence over the specificity of what information is prioritized in memory, rather than a general enhancement of all memory, and provide insight into how BLA-MTL projections contribute to the dynamics of memory prioritization.

## Introduction

Decades of learning and memory research suggest the basolateral amygdala (BLA) plays an essential role in enhancing emotional memories^1–11^. However, there is uncertainty as to whether this enhancement is a generalized boost or prioritizes some information over others. Neuroscience studies using direct perturbations suggest the basolateral amygdala (BLA) mediates a causal influence on many different types of memory more generally^12–15^. In contrast, psychology-inspired studies using indirect and correlational approaches propose that specificity exists for how the BLA prioritizes experiences in memory^16,17^. A limitation of this prior work is that numerous variables, including attention and arousal, covary with putative engagement of the BLA, making interpretation of the findings difficult. To circumvent these challenges, some studies in both rodents and humans have focused on direct manipulations of the BLA in the post-training memory consolidation period to dissociate learning effects from performance effects. However, to our knowledge, no prior study has used direct manipulations in this manner to investigate whether the human BLA prioritizes certain kinds of memories over others.

Increasing evidence in both humans and rodents suggests that BLA modulation of memory likely occurs via BLA interactions with other brain regions, including structures in the medial temporal lobe (MTL)^18–21^. Furthermore, research on human memory and direct perturbations of the BLA in rodents demonstrate the potential of BLA stimulation to interrogate these downstream connections^2–6,12,19–28^. For example, prior work in rodents suggests that memory-enhancing electrical stimulation of the BLA induces theta-modulated gamma oscillations in the hippocampus^29^, an oscillatory state previously correlated with good memory^30,31^. Similarly, recent work in humans found that direct electrical stimulation of the BLA enhances declarative memory, and this memory enhancement is marked by interactions between the BLA and hippocampus^20^. Together, these results from both rodent and human studies show that BLA modulation can induce specific memory-related signals in downstream regions. More specifically, direct activation of the BLA can enhance memory by modulating plasticity in the hippocampus.

Prior neuroanatomy studies using fluorescent tract tracing in both rodents and nonhuman primates suggest that BLA inputs to the hippocampus and adjacent MTL structures exhibit a pattern that approximates an anterior-to-posterior gradient of stronger to weaker projections^32,33^. Specifically, the BLA sends more dense excitatory projections to the anterior hippocampus (aHC; the homolog of ventral hippocampus in rodents) and perirhinal cortex compared to the posterior hippocampus (pHC; dorsal hippocampus in rodents) and parahippocampal cortex^32,33^. Evidence from rats, humans, and non-human primates suggests that this anterior-to-posterior pattern of BLA connectivity parallels a functional trend of separation of non-spatial object information from that about spatial contexts^34–37^. Consequently, the BLA appears to be anatomically situated to modulate memory for objects more so than memory for discrete scenes or spatial contexts. Nevertheless, prior studies have not yet investigated whether and how the BLA modulates memory for multiple types of memory in the same the experiment. Thus, the premise that the BLA prioritizes some kinds of memories over others has not been directly tested.

Movement towards novel therapeutics designed to emulate endogenous mechanisms of memory enhancement requires an understanding of the basic science of normal memory function. The present study therefore sought to test whether directly engaging the amygdala results in a generalized modulation of memory or differentially prioritizes memory processes for objects versus scenes – types of information processed by distinct memory systems – through a more specific influence on memory. We tested whether brief electrical stimulation applied directly to the human BLA could enhance recognition memory for specific images of neutral object or scenes without eliciting an emotional experience. Eight patients with depth electrodes placed in the amygdala, hippocampus, and other frontal-temporal lobe targets for clinical purposes performed a recognition-memory task for object and scene images. During encoding, half of the images were immediately followed by a 1-s low-amplitude (0.5-1 mA) theta-modulated gamma frequency (theta-burst) electrical stimulation to the amygdala, and one day later participants were given a retrieval test. Amygdala stimulation at this low amplitude reliably improved later object-recognition memory without eliciting emotional responses and had no effect on scene-recognition memory. These results highlight the utility of stimulation to initiate endogenous memory prioritization processes in the absence of emotional input and to our knowledge are the first to suggest that stimulation of the human amygdala has the capacity to prioritize enhancement of specific declarative memories over others rather than a general enhancement of all memory.

To investigate the neural substrates underlying BLA stimulation and the prioritization of object memory, we also considered the relationship between the degree of BLA-hippocampal connectivity and the resulting memory effect. Effective connectivity can be used to determine the causal influence between regions in the human brain^38–41^. Recent work using effective connectivity to map human brain networks shows that single-pulse intracranial stimulation triggers a local electrical response at the stimulation site, as well as evoked responses (single pulse evoked potentials; SPEP) at downstream locations, in proportion to the strength of the effective connection between the two locations^42,43^. The magnitude of the SPEP provides a measure of connectivity that is correlated with both structural^44^ and functional^45,46^ connectivity among brain regions, with sufficient resolution to infer the direction of information flow between two regions^38^. Importantly, where theta-burst stimulation is useful for modulating memory, it is an ineffective stimulation pattern for probing network connectivity. Therefore, we investigated whether and how single-pulse electrical manipulations of the human BLA differentially influence the amplitude of evoked responses in the aHC and pHC as a measure of BLA-MTL connectivity in the same participants who completed the objects and scenes recognition memory task. We found that electrical stimulation of the BLA reveals stronger effective connectivity with the aHC compared to the pHC. These results provide a potential circuit mechanism by which the amygdala prioritizes object memory over scene memory in humans. Our findings reconcile the neuroscience and psychology literature to suggest the BLA mediates a more specific memory enhancement rather than a generalizable influence on memory, at least in part, due to its projections to the MTL.

## Materials and Methods

### Participants

Eight participants (five females, three males) with drug-resistant epilepsy volunteered and provided written informed consent for all study procedures (see Supplemental material for inclusion/exclusion criteria). The study protocol was approved by both the University of Utah and Washington University St. Louis Institutional Review Boards. Stereotactic depth electrodes were implanted into the brain parenchyma by neurosurgeons for the sole purpose of clinical seizure investigation.

### Brain Stimulation and Recording

Electrode contacts were localized using LeGUI^47^ or VERA^48^, semi-automated detection and localization platforms for intracranial electrodes, following coregistration of pre- and postoperative brain imaging (Fig. 1A). Theta-burst stimulation parameters replicated those used in prior rat^19,21,49^ and a human study^20^ that demonstrated amygdala-mediated memory enhancement and physiological neuromodulation. Specifically, bipolar stimulation was delivered unilaterally to the BLA in current-regulated, charge-balanced, cathode-first biphasic rectangular pulses at 1.0 mA (or 0.5 mA for three of the 14 sessions) for 1 s in eight trains of four pulses at 50 Hz (Fig. 1C). Single-pulse bipolar stimulation for the effective connectivity analyses was applied to the BLA at 3.0 mA with a 1-3 s interpulse interval (Fig. 4A; See Table S3 for patient-specific stimulation locations). SPEPs were recorded at a sampling rate of 2,000 Hz. Stimulation did not elicit seizure activity or afterdischarges in a thorough pre-test stimulation safety session conducted with a clinical epileptologist for each patient.

**Figure 1.**
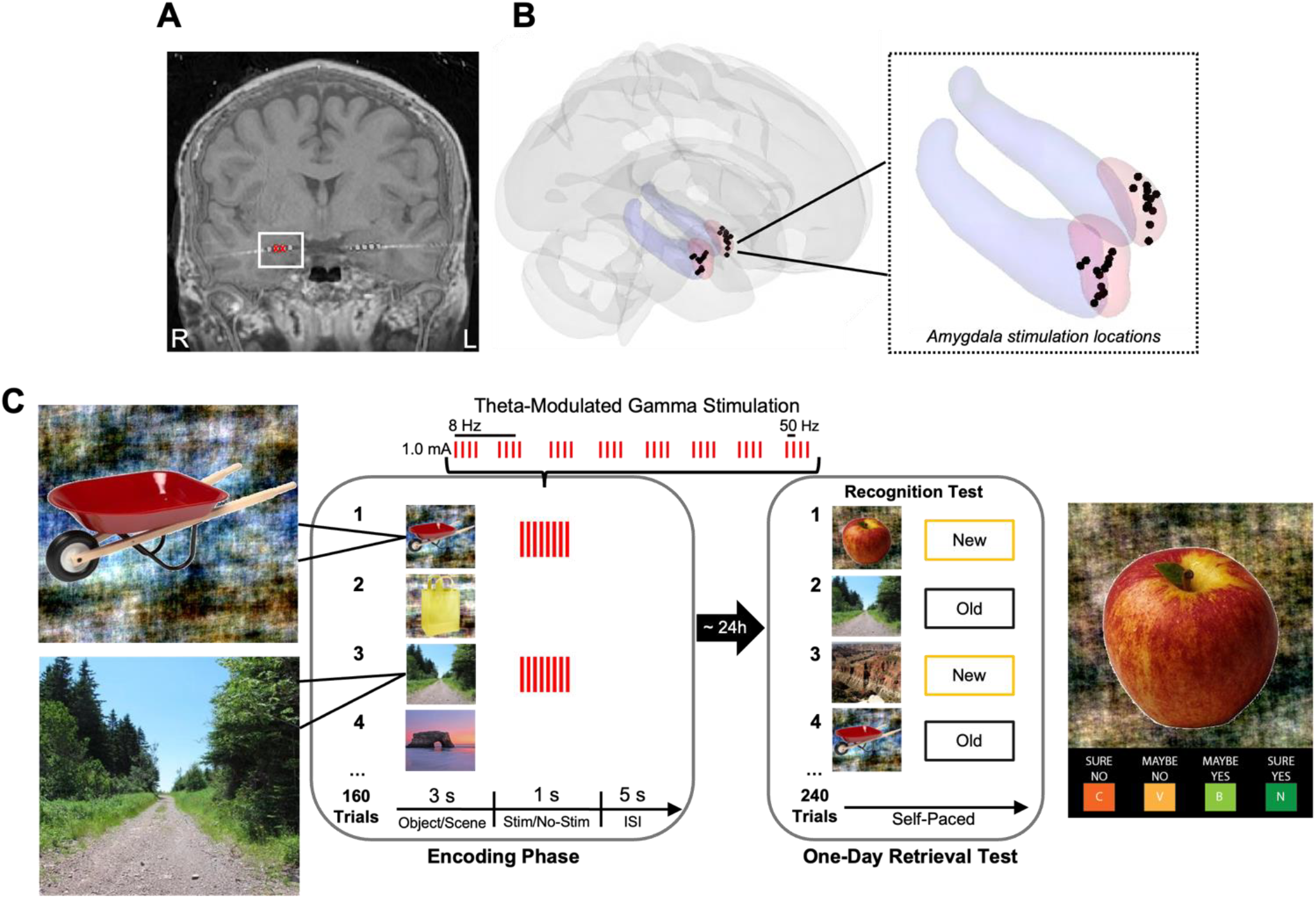
The procedure used to stimulate the human amygdala and to test recognition memory. **A**, A representative postoperative coronal MRI/CT showing electrode contacts in the BLA (white square). The red Xs denote the bipolar pair of contacts stimulated in this example. **B**, Brain template in Montreal Neurological Institute (MNI) space on which all amygdala stimulation electrodes have been plotted. The amygdala is highlighted in red, and the hippocampus is plotted in blue for reference. **C**, Schematic of the recognition-memory task in which the BLA was stimulated (1-s stimulation pulse sequence; each pulse = 500 *μ*s biphasic square wave; pulse frequency = 50 Hz; train frequency = 8 Hz) after the presentation of half of the images in the encoding phase and recognition memory was tested for all images one day after the encoding phase. Example object and scene images (left). The object images were presented on scrambled backgrounds that were created to preserve the overall luminance, hue, spatial frequency, and shape between object and scene images (see Supplemental material for more details on the stimuli). Example retrieval test image with the response options presented for each image (right).

### Behavioral Tasks

#### Recognition-memory task procedure

During the encoding phase, each participant viewed 160 images, 80 neutral object images, and 80 outdoor scene images, one trial at a time (Fig. 1C) (see Supplemental material for additional details on the stimuli). Participants were asked to make a “like” or “dislike” judgment with respect to the image on each trial (or in the case of three of the 14 sessions a “memorable” or “not memorable” judgment) to orient and encourage participants to pay attention to each image. A fixation screen was presented for a jittered duration of 0.5-1.5 s immediately before each image onset. Each image was presented for 3 s, and for a randomly selected half of the trials (stimulation condition), an offset of the image coincided with the onset of 1 s of stimulation to the BLA. The interstimulus interval was 5 s. Memory was tested approximately one day later (mean = 22 h, range = 18-25 h delay) via a self-paced old/new recognition-memory test for all 160 images. Eighty new images of each type served as foils (new images only occurred during the one-day retrieval test and were never associated with stimulation). Foil images were used to assess each participant’s propensity to guess (false alarm) and performance on these new images was used as an exclusion criterion for the experiment (see Supplemental material for additional details on exclusion criteria). After making old/new judgments, participants also made sure/maybe certainty judgments. No electrical stimulation was delivered during the memory retrieval test. Some participants completed the memory task multiple times on different days with a new image set and a different amygdala stimulation location. Thus, across the eight participants, a total of 14 different testing sessions were conducted. (Linear mixed effects modeling was used to account for the multiple testing sessions within each patient. See Supplemental material for additional details on the analysis procedure.)

#### Stimulation awareness test procedure

After the one-day retrieval test, participants were administered 20 trials in which stimulation or sham stimulation (no current) was randomly delivered to the BLA (Fig. 3A). Participants were asked “Did you feel anything?” Stimulation to the BLA using the same parameters as during the memory task was delivered, and participants were blind to which trials were paired with stimulation.

### Electrophysiological Data Analysis

Analyses of electrophysiological data focused on evoked potentials recorded from the hippocampus in five of the eight total patients who had both anterior and posterior hippocampal electrode sampling for seven different BLA stimulation locations. To analyze the SPEPs for the aHC and pHC we created peri-stimulation epochs for each stimulation trial per region and voltages were normalized (z-scored) to the prestimulation baseline data segment per stimulation trial. We measured the normalized peak amplitude of the SPEP in the 100-ms poststimulation period^50^ and calculated the trial-averaged voltage peak amplitude. To control for individual differences, we compared the SPEP amplitude in the aHC to the SPEP amplitude in the pHC for each patient/stimulation location.

## Results

The influence of amygdala stimulation on recognition-memory performance for neutral object and scene images was assessed in eight human participants across 14 different memory encoding/retrieval sessions (see Fig. 1, Table S3, and Table S1 for information on electrode locations and patient demographics). Fig. 1C illustrates the recognition-memory procedure and depicts the 1 s of low-amplitude electrical stimulation (eight trains of 50 Hz, 0.5-1.0 mA biphasic pulses) that was delivered to the amygdala at the offset of image presentation for a randomly selected half of the object and scene images during the encoding phase of the task. Memory performance was tested one day later. Fig. 1A shows a representative postoperative coronal CT/MRI depicting electrode contacts in the amygdala for one patient. Fig. 1B represents all bipolar amygdala stimulation locations across all subjects plotted in MNI space.

### Electrical Stimulation of the BLA Enhances Object Memory but Not Scene Memory

Fig. 2A shows the memory outcomes as a standard discriminability index (d’), a typical approach for representing memory performance on a recognition task that accounts for an individual’s propensity to guess (for individual Hit and False Alarm rates see Fig. S4). Results indicate that brief stimulation of the BLA during the encoding phase enhanced object-recognition memory relative to control (no-stimulation) object images the next day [F_(1,26)_ = 24.67, *p* < 0.0001]. By comparison, stimulation did not affect scene-recognition memory performance [F_(1,26)_ = 0.28, *p* = 0.60]. Although a standard 2 × 2 model did not yield a significant interaction for objects and scenes [F_(1,52)_ = 1.63, *p* = 0.21], we next analyzed more explicitly whether the magnitude of the amygdala-mediated memory enhancement was greater for objects versus scenes. Fig. 2B shows the recognition-memory performance plotted for each session as the difference in d’ between the stimulation and no-stimulation conditions for objects and scenes. Object images had a higher d’ difference score on average compared to scene images [objects vs. scenes d′ difference: mean(SEM) = 0.23(0.048) vs. −0.040(0.077), F_(1,26)_ = 11.45, *p* = 0.0023]. (See Table S4 for patient-specific recognition-memory scores.)

**Figure 2.**
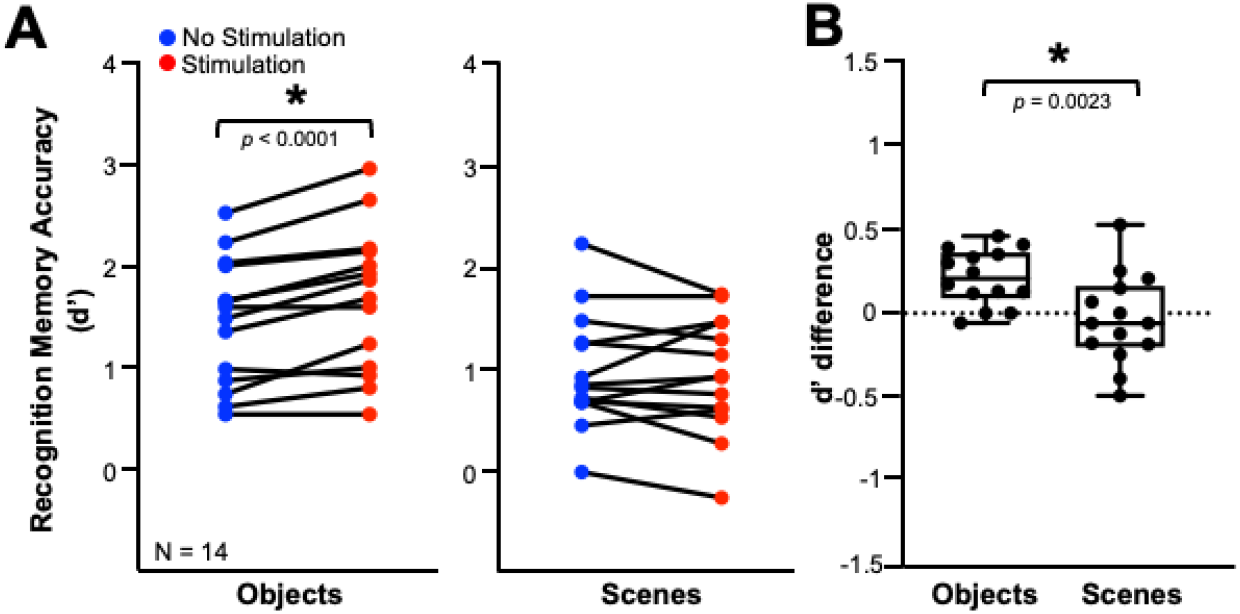
Brief electrical stimulation of the BLA in humans enhanced subsequent memory for objects but not scenes. **A**, Recognition-memory performance for each of the 14 sessions across eight patients plotted as discriminability index (d’) for object and scene images. **B**, Recognition-memory performance plotted for each session as the difference in d’ in the stimulation and no-stimulation conditions (scatter plots in A). Overlaid box-and-whisker plots show the median, range, and interquartile range for each condition. \**p* < 0.05 compared with control images.

### BLA Stimulation Does Not Influence Memory Confidence

In the present recognition-memory task patients were not only asked to mark images as ‘old’ or ‘new’, but also how certain they were in their response and whether they had a conscious recollection of the images (‘sure’) or simply know they had been encountered before on some other basis (‘maybe’). Fig. S3 represents the analysis of these certainty judgments and shows that patients indicated they were “sure” more often for stimulated versus nonstimulated object images [Fig. S3A; F_(1,26)_ = 4.26, *p* = 0.049]. However, patients also indicated they had “maybe” encountered an image before more often for stimulated versus nonstimulated objects [Fig. S3C; F_(1,26)_ = 8.90, *p* = 0.0061]. This implies our main effect in Fig. 2 is the same whether we restrict our analyses to certain or uncertain responses, suggesting BLA stimulation enhances overall memory strength.

### BLA Stimulation Enhances Memory in the Absence of Subjective Awareness

Fig. 3 shows the results of a subsequent stimulation awareness test in which participants were asked to indicate whether amygdala stimulation had been delivered across 10 BLA-stimulation and 10 sham-stimulation trials that were presented in a random order. Patients’ responses indicate they were unable to reliably discriminate if or when any stimulation was being delivered (Fig. 3C) [*t*_(11)_ = 1.24, *p* = 0.24]. Furthermore, no patient reported emotional responses associated with the 80 stimulation trials during the memory encoding phase of the recognition-memory task. Thus, the amygdala stimulation used in the present study was capable of enhancing one-day object recognition memory in the absence of any subjective emotional response.

**Figure 3.**
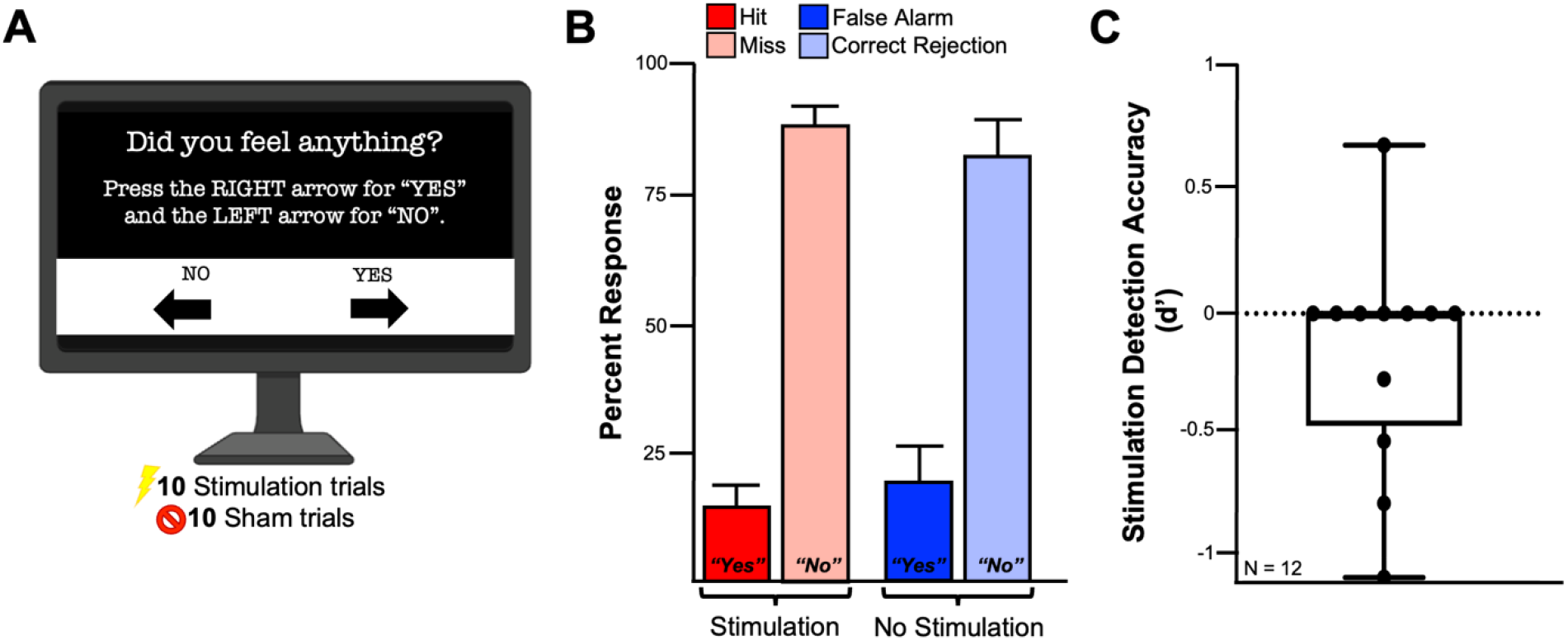
Participants could not reliably detect the presence of amygdala stimulation. **A**, Illustration of the stimulation awareness test. Patients were given 10 amygdala stimulation trials and 10 sham/no-stimulation trials and were asked to report any sensations following each trial. Patients were blind to whether the trials were stimulation or sham trials. **B**, Reported responses of patients when they were asked in stimulation awareness testing whether they felt any sensation of stimulation. Hits (correct “Yes” response for a stimulation trial), Misses (incorrect “No” response for a stimulation trial), False Alarms (incorrect “Yes” response for a no-stimulation trial), and Correct Rejections (correct “No” response for a no-stimulation trial). **C**, Stimulation detection ability for each patient plotted as discriminability index (d’), incorporating the normalized hit and false alarm rates from panel B. The one patient that appears to reliably detect the stimulation (with a d’ value of 0.62) had a Hit rate of 30%, False Alarm rate of 10%, Miss rate of 70%, and Correct Rejection rate of 90%.

### BLA Stimulation Does Not Influence Reaction Time

No differences in reaction times during the one-day retrieval test were observed between stimulation and no-stimulation control images, suggesting that amygdala-mediated memory enhancement did not produce reduced reaction times in subsequent testing (Fig. S2B). However, irrespective of stimulation and no-stimulation condition, reaction times were significantly longer for scene images compared to object images during the one-day retrieval test (Fig. S2A) [scenes vs. objects reaction time: mean(SEM) = 2.84(0.23) vs. 2.22(0.14), t (27) = 4.76, *p* < 0.0001].

### Single-Pulse Electrical Stimulation Reveals Stronger Effective Connectivity Between the BLA and aHC Compared to the BLA and pHC

Fig. 4A illustrates the stimulation procedure used to examine effective connectivity between the BLA and the hippocampus. High amplitude (3 mA) single-pulse electrical stimulation was delivered to the BLA while evoked responses were recorded in the aHC and pHC. (No patient reported feeling the presence of this stimulation.) Fig. 4B shows a representative example of a trial-averaged waveform in the aHC and pHC for one patient. Fig. 4C shows the trial-averaged voltage peak amplitudes of the stimulation-evoked SPEPs averaged across all aHC electrode contacts and pHC contacts for each stimulation session. Fig. 4D shows the SPEP amplitudes plotted for each session as the difference in amplitude between the aHC and pHC. Stimulation revealed significantly stronger effective connectivity, as measured by the magnitude of the evoked responses, in the aHC networks important for object memory compared to the pHC networks important for scene memory [F_(1,12)_ = 5.87, *p* = 0.032]. Together, these results suggest stronger effective connectivity between the BLA and aHC compared to the BLA and pHC, providing a potential mechanism by which BLA stimulation enhances object memory over scene memory.

**Figure 4.**
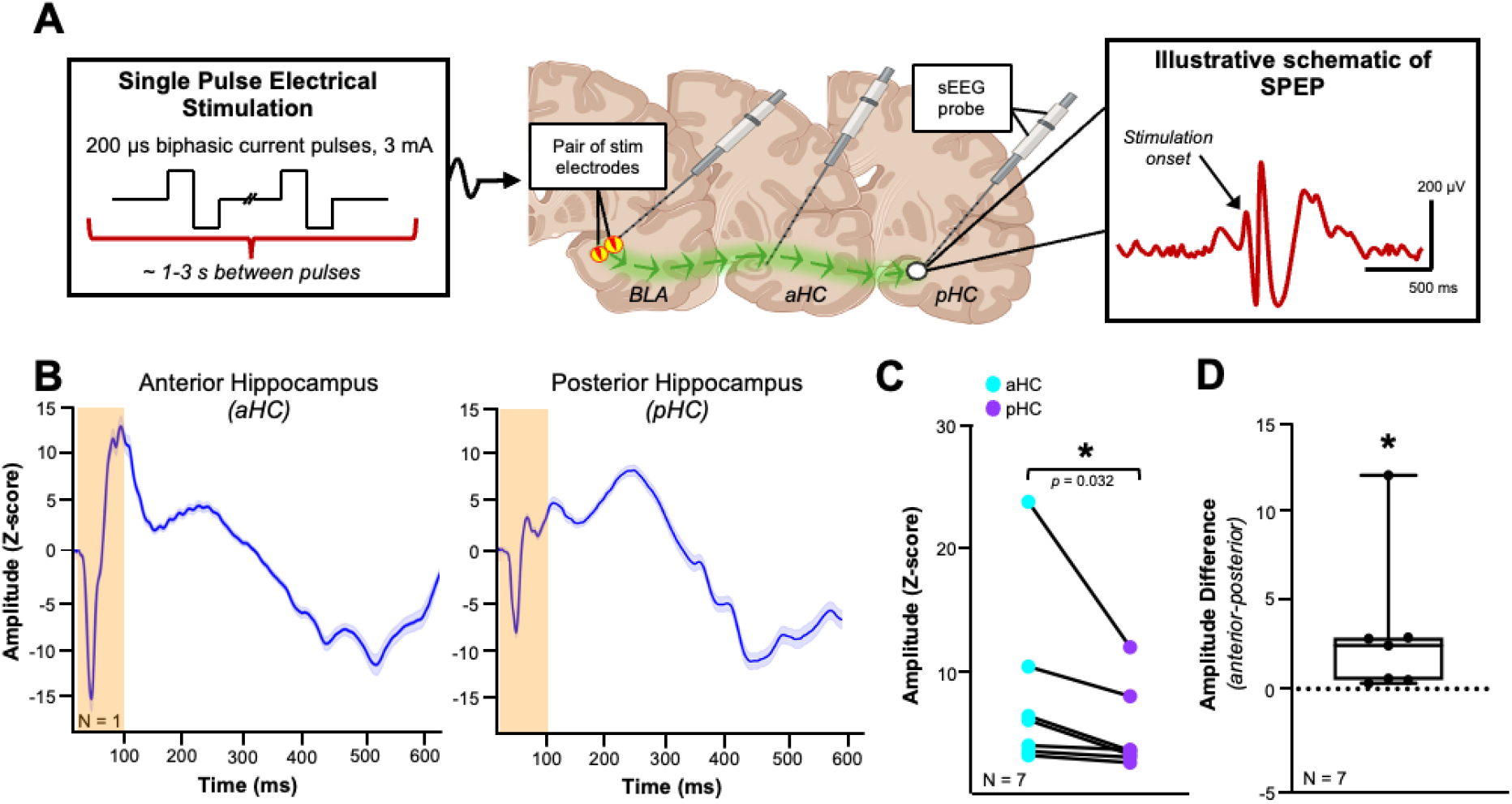
Single-pulse electrical stimulation of the BLA induces higher magnitude responses in the anterior compared to posterior hippocampus. **A**, Schematic of the stimulation procedure used to deliver high amplitude single-pulse stimulation to the BLA, and record evoked potential responses in the hippocampus (created with BioRender.com). **B**, Example of a trial-averaged single-pulse evoked potential (SPEP) recorded from one anterior hippocampus channel (left) and one posterior hippocampus channel (right) from one patient. Stimulation occurred at time = 0 ms. Shading around each SPEP trace represents the SEM for the 30 trials of stimulation delivered. The orange shaded area represents the time window used to extract the amplitude. **C**, The maximum stimulation response amplitude recorded in the 100 ms poststimulation averaged across all anterior and posterior hippocampus electrodes for a subset of patients/stimulation locations included in the recognition-memory procedure. **D**, The SPEP amplitudes plotted as the difference between the anterior and posterior responses (scatter plots in C). \**p* < 0.05.

## Discussion

The present study aimed to determine whether neuroanatomy-inspired direct manipulation of the BLA would reveal the psychological specificity of the memory enhancement for different types of visual stimuli. Our results suggest that direct electrical stimulation of the human amygdala reliably improves long-term recognition memory for images of neutral objects, but not for scenes, types of information processed by distinct memory systems in the anterior and posterior medial temporal lobe (MTL), respectively. Furthermore, this memory enhancement occurred in the absence of a subjective emotional response, indicating a causal role for the amygdala in memory that is dissociable from emotional experience. Finally, our results provide evidence for stronger effective connectivity between the BLA and aHC compared to the BLA and pHC, reconciling the neuroscience and psychology literature to suggest the BLA mediates a more specific memory enhancement rather than a generalizable influence on memory, at least in part, due to its projections to the MTL.

### Temporally Specific Memory Enhancement

Stimulation of the BLA enhanced object memory in an “event” specific (trial-specific) manner. Indeed, with amygdala-mediated memory enhancement, we not only observe stimulus specificity for object memory but also temporally specific learning that is robust after a single trial and persists the following day. Specifically, no carryover effects of stimulation on memory were observed for control images that followed stimulation trials during the study phase (Fig. S1). The lack of carryover indicates that the object memory enhancement was temporally specific. That is, objects in the stimulation condition were remembered better than control objects initially viewed only seconds (~ 7 s) apart. This temporal specificity implies that long-lasting effects such as increases in blood hormone levels are unlikely to explain the observed memory enhancement. Furthermore, this temporal specificity is consistent with evidence that BLA activity in the consolidation period immediately following an experience is important for later retrieval of that experience^12^. A possible mechanism for the observed temporally specific amygdala-mediated memory enhancement involves changes in synaptic strength following an experience. Evidence suggests change in synapse function is involved in memory storage, and that the BLA in particular modulates the molecular processes that occur during the consolidation period^51^. Prior work shows that the BLA modulates hippocampal synaptic plasticity and modulates the expression of plasticity-associated proteins like ARC in a memory-dependent fashion^52,53^. Furthermore, memory-influencing BLA stimulation enhances ARC levels in the hippocampus^53^, providing a possible molecular mechanism by which our direct electrical stimulation of the BLA enhances object memory in a temporally precise manner.

### Mechanisms of Amygdala-Mediated Memory Prioritization

The present study suggests that a potential circuit mechanism by which BLA stimulation prioritizes object memory over scene memory involves increased effective connectivity between the BLA and aHC, compared to the BLA and pHC. Specifically, single-pulse electrical stimulation of the BLA resulted in higher amplitude SPEPs in the aHC versus pHC which is consistent with our finding that direct electrical stimulation of the BLA enhances object memory but not scene memory. These results suggest the BLA influences the specificity of what information is prioritized in memory rather than a general enhancement of all memory, and that BLA interactions with the hippocampus may be important for this specificity.

A historical tension exists between the neuroscience and psychology literature, such that neuroscience studies using direct perturbations suggests the BLA wields a generalizable influence on memory^12–15^. In contrast, emotional memory studies from the psychology literature, using more indirect interrogations, propose that specificity exists for which experiences are prioritized in memory^16,17^. To our knowledge, no prior studies exist in which direct manipulations of the BLA were shown to improve one type of memory but not another. Critically, these studies did not investigate multiple memory types in the same the experiment. In contrast, the psychology literature is replete with examples of how emotion enhances memory in some instances and impairs or has no effect on memory in other cases^16,54^. One possible explanation for the inconsistencies in the psychology literature is the imprecision of the experimental manipulations available to researchers, combined with the numerous indirect routes by which emotion can influence memory. However, the present study reconciles the neuroscience and psychology literatures, suggesting that BLA stimulation has the capacity to influence certain memories over others in the absence of emotional response. The projections from the BLA to the hippocampus are one potential source for how certain memories are prioritized over others.

### Future Investigations

To further understand the BLA’s prioritization of object learning, it is also critical to investigate the direct relationship between the areas stimulated and the resulting behavioral effect. Neurostimulation modeling is an increasingly popular tool for predicting the effects of stimulation on neural tissue in the human brain and relating these effects to meaningful behavioral changes^55^. One aspect of neurostimulation modeling involves examining the volume of neural tissue activated (VTA)^56^. VTA estimates incorporate electric field distribution to quantify the location and extent of neuronal activation caused during electrical stimulation^55^. Furthermore, recent developments in neurostimulation modeling approaches integrate DTI neuroimaging data to examine the underlying white matter anatomy and structural connectivity to estimate the effects of electrical stimulation directly on the prevalent tracts^57,58^. Numerous clinical studies have recently implemented neurostimulation modeling techniques in the human brain to map the local and network effects of deep brain stimulation (DBS)^59–61^. For example, DBS targeting the fornix may enhance cognitive function in patients with Alzheimer’s disease and neurostimulation modeling approaches suggest that variable outcome is correlated to the effects of focal electric fields on specific white matter tracts traversing the stimulation volumes^59^. Thus, neurostimulation modeling is a powerful approach for examining the intersection between the VTA and estimated neural pathway activation in studies that use direct electrical stimulation of the human brain. Future work will investigate whether and how stimulation of the human BLA differentially activates BLA-aHC and BLA-pHC structural connections and does so in a manner related to memory for objects, scenes, and other types of stimuli.

### Additional Considerations

Our study raises several methodological and interpretive considerations. First, as in any study examining the effects of direct electrical stimulation in subjects undergoing intracranial monitoring, our study necessarily consisted of patients with epilepsy rather than healthy participants. Intracranial monitoring samples extensive territories of the brain representing normal and pathological tissue, to define the boundaries of each patient’s seizure focus accurately^62,63^. Although specific areas of each patient’s brain were presumed to be functioning abnormally, we accounted for the clinical heterogeneity across patients by using a within-subject design, comparing each patient’s memory performance in the stimulated versus nonstimulated conditions. Memory-enhancement effects were observed regardless of the differences in seizure foci across patients, including several subjects in which the hippocampus ipsilateral to BLA stimulation was implicated in the seizure focus (Table S2). Comparable memory-enhancement results resembled those from prior work in humans^20^ and several prior studies in rats without seizures^19,21,49^, suggesting that our findings are unrelated to epilepsy. Second, the present findings observed variability in the baseline memory performance for objects versus scenes. Specifically, the recognition-memory accuracy for nonstimulated scene images was lower than that of nonstimulated object images [F_(1,26)_ = 6.06, *p* = 0.021; data not shown], likely due in part to the differences in nameability^64^ between objects and scenes, raising the possibility that the lack of memory enhancement for scene images was due to a floor/ceiling effect. Although based on the numerical averages for recognition-memory accuracy, there was still sufficient statistical room for a scene memory enhancement to be observed. Moreover, prior learning and memory studies suggest that weaker baseline memory performance tends to show the largest stimulation-mediated memory improvement^65,20^. Nonetheless, it is difficult to be certain that a neurobiological floor effect, in which the memory itself could not be enhanced, did not occur. Finally, the observed memory-enhancement effect for objects was reliable and significant with no effect on scene memory, in contrast to previous studies that implicated the BLA in spatial and contextual learning. Evidence in rodents suggests optogenetic stimulation of the BLA enhances spatial memory^18^. However, this prior work targeted BLA projections to the medial entorhinal cortex, suggesting that pathway manipulations are important for BLA modulation of spatial memory. Although we are unable to deliver electrical stimulation to distinct BLA projections in the human brain, our stimulation more fully engages the widespread projections of the BLA, and our results suggest stimulation preferentially engages the denser anatomical projections from the BLA to the aHC.

## Conclusions

We leveraged electrical stimulation to determine what one can uniquely learn from direct manipulation of the BLA in the human brain. Our results suggest BLA stimulation elicits no subjective emotional response but enhances long-term memory for object images but not scene images. Furthermore, single pulse electrical stimulation of the BLA elicits higher magnitude evoked responses in the aHC compared to the pHC, suggesting a potential mechanism by which BLA stimulation enhances object memory but not scene memory. Our findings reconcile the neuroscience and psychology literature to suggest the BLA mediates an important influence over the specificity of what information is prioritized in memory, rather than a general enhancement of all memory, and that BLA interactions with the hippocampus may be important for this specificity. These experiments provide critical knowledge regarding how human MTL network connectivity, rather than individual brain structures, influences memory consolidation. Moreover, these findings represent a significant advancement in the basic science of normal memory function and a step towards novel therapeutics designed to emulate endogenous mechanisms of amygdala-mediated memory enhancement.

## Resource Availability

### Lead contact

Further information and requests for resources should be directed to and will be fulfilled by the lead contact, Krista L. Wahlstrom

### Materials availability

This study did not generate new unique reagents.

### Data and code availability

The datasets generated are available through the NIMH Data Archive and the code used to analyze the current study’s data are available on the INMAN Laboratory’s GitHub repository.

## Acknowledgements

We thank the patients, physicians, and staff of the University of Utah and Washington University St. Louis Epilepsy Monitoring Units for their time and trust in this work. Efforts and resources for this project were supported by NIH Grant R01MH120194 (to C.S.I., J.T.W., and K.L.W.)

## CRediT Author Statement

**Krista Wahlstrom**: Conceptualization, Methodology, Software, Formal Analysis, Investigation, Data Curation, Project Administration, Writing – Original Draft, Writing – Review & Editing, Visualization; **Justin Campbell**: Investigation, Formal Analysis, Writing – Review & Editing; **Martina Hollearn**: Investigation, Writing – Review & Editing; **James Swift**: Software, Investigation, Formal Analysis, Writing – Review & Editing; **Markus Adamek**: Software, Investigation, Writing – Review & Editing; **Lou Blanpain**: Methodology, Writing – Review & Editing; **Tao Xie**: Investigation, Writing – Review & Editing; **Peter Brunner**: Software, Investigation, Supervision, Writing – Review & Editing; **Stephan Hamann**: Conceptualization, Funding Acquisition, Writing – Review & Editing; **Amir Arain**: Resources, Writing – Review & Editing; **Lawrence Eisenman**: Resources, Writing – Review & Editing; **John Rolston**: Resources, Writing – Review & Editing; **Shervin Rahimpour**: Resources, Writing – Review & Editing; **Joseph Manns**: Conceptualization, Supervision, Project Administration, Funding Acquisition, Writing – Review & Editing; **Jon Willie**: Conceptualization, Investigation, Supervision, Resources, Project Administration, Funding Acquisition, Writing – Review & Editing. **Cory Inman**: Conceptualization, Supervision, Resources, Project Administration, Funding Acquisition, Writing – Review & Editing.

## Declaration of Interests

The authors declare no competing interests.

## Additional Resources

This study was conducted as part of a National Institutes of Health clinical trial (NCT05065450).

## Supplemental Information

Document S1. Figures S1-S4 and Tables S1-S4

## Supplemental Figure Legends

**Figure S1. Recognition-memory performance for objects followed by stimulation relative to no-stimulation objects that came after stimulation. A**, Recognition-memory performance (stimulation trials – no-stimulation trials after stimulation trials) for each session plotted as discriminability index (d’) for objects. **B**, Recognition-memory performance plotted for each session as the difference in d’ between the stimulation and no-stimulation-after-stimulation objects. If stimulation enhanced memory for subsequent no-stimulation images, then the data would show a decrease of the original memory-enhancement effect across sessions, but if stimulation diminished memory for subsequent no-stimulation images, then the data would show an increase of the original memory-enhancement effect across sessions. These results suggest no evidence of a change in our original effect and thus no significant carry-over effect of stimulation trials upon the subsequent encoding trials. \**p* < 0.05.

**Figure S2. Reaction times (RT) for recognition of previously viewed and new images at the one-day retrieval test. A**, Mean RTs across all retrieval sessions for previously viewed object and scene images. **B**, Mean RTs across all retrieval sessions for previously nonstimulated, previously stimulated, and new images for objects *(left)* and scenes *(right)*. **C**, Mean RTs across all retrieval sessions for Hits (correct “old” response for an old image), Misses (incorrect “new” response for an old image), False Alarms (incorrect “old” response for a new image), and Correct Rejections (correct “new” response for a new image). **D**, Mean RTs across all retrieval sessions for Hits, Misses, False Alarms, and Correct Rejections for objects *(left)* and scenes *(right)*. **E**, Mean RTs across all retrieval sessions for previously nonstimulated Hits, previously stimulated Hits, previously nonstimulated Misses, and previously stimulated Misses. \**p* < 0.05.

**Figure S3. BLA stimulation does not influence memory confidence. A**, Recognition-memory performance for each session plotted as discriminability index (d’) for object and scene images for “Sure” responses. **B**, Recognition-memory performance plotted for each session as the difference in d’ in the stimulation and no-stimulation conditions for the “Sure” responses (scatter plots in C). Overlaid box-and-whisker plots show the median, range, and interquartile range for each condition. **C**, Recognition-memory performance for each session plotted as discriminability index (d’) for object and scene images for “Maybe” responses. **D**, Recognition-memory performance plotted for each session as the difference in d’ in the stimulation and no-stimulation conditions for the “Maybe” responses (scatter plots in A). Overlaid box-and-whisker plots show the median, range, and interquartile range for each condition. \**p* < 0.05 compared with control images.

**Figure S4. Recognition memory performance for previously viewed images and new images during retrieval testing. A**, Number of responses in percent of Hits (correct “old” response for an old image) relative to the total number of old images for previously stimulated and nonstimulated object *(left)* and scene *(right)* images. **B**, Number of confident/”Sure” responses in percent of Hits relative to the total number of “Sure” responses for previously stimulated and nonstimulated object *(left)* and scene *(right)* images. This panel suggests the main results in panel A are essentially the same when we restrict the analyses to the putatively highest-quality data. **C**, Number of responses in percent of False Alarms (incorrect “old” response for a new image) relative to the total number of new images presented at retrieval testing *(left)*, and number of confident/”Sure” responses in percent of False Alarms relative to the total number of “Sure” responses *(right)*. \**p* < 0.05; &*p* < 0.10.

## Notes

### Competing Interest Statement

The authors have declared no competing interest.

## References

1. Zheng, J., Anderson, K.L., Leal, S.L., Shestyuk, A., Gulsen, G., Mnatsakanyan, L., Vadera, S., Hsu, F.P.K., Yassa, M.A., Knight, R.T., et al. (2017). Amygdala-hippocampal dynamics during salient information processing. Nature Communications 8, ncomms14413. 10.1038/ncomms14413.

2. Richardson, M.P., Strange, B.A., and Dolan, R.J. (2004). Encoding of emotional memories depends on amygdala and hippocampus and their interactions. Nat Neurosci 7, 278–285. 10.1038/nn1190.

3. McGaugh, J.L. (2013). Making lasting memories: Remembering the significant. Proc Natl Acad Sci U S A 110, 10402–10407. 10.1073/pnas.1301209110.

4. Cahill, L., Babinsky, R., Markowitsch, H.J., and McGaugh, J.L. (1995). The amygdala and emotional memory. Nature 377, 295–296. 10.1038/377295a0.

5. Cahill, L., Haier, R.J., Fallon, J., Alkire, M.T., Tang, C., Keator, D., Wu, J., and McGaugh, J.L. (1996). Amygdala activity at encoding correlated with long-term, free recall of emotional information. Proc Natl Acad Sci U S A 93, 8016–8021.

6. Hamann, S. (2001). Cognitive and neural mechanisms of emotional memory. Trends in Cognitive Sciences 5, 394–400. 10.1016/S1364-6613(00)01707-1.

7. Phelps, E.A. (2004). Human emotion and memory: interactions of the amygdala and hippocampal complex. Current Opinion in Neurobiology 14, 198–202. 10.1016/j.conb.2004.03.015.

8. LaBar, K.S., and Cabeza, R. (2006). Cognitive neuroscience of emotional memory. Nat Rev Neurosci 7, 54–64. 10.1038/nrn1825.

9. Kilpatrick, L. (2003). Amygdala modulation of parahippocampal and frontal regions during emotionally influenced memory storage. NeuroImage 20, 2091–2099. 10.1016/j.neuroimage.2003.08.006.

10. Dolcos, F., LaBar, K.S., and Cabeza, R. (2004). Interaction between the Amygdala and the Medial Temporal Lobe Memory System Predicts Better Memory for Emotional Events. Neuron 42, 855–863. 10.1016/S0896-6273(04)00289-2.

11. Canli, T., Zhao, Z., Brewer, J., Gabrieli, J.D.E., and Cahill, L. (2000). Event-Related Activation in the Human Amygdala Associates with Later Memory for Individual Emotional Experience. J. Neurosci. 20, RC99–RC99.

12. McGaugh, J.L. (2004). The amygdala modulates the consolidation of memories of emotionally arousing experiences. Annu Rev Neurosci 27, 1–28. 10.1146/annurev.neuro.27.070203.144157.

13. Huff, M.L., Miller, R.L., Deisseroth, K., Moorman, D.E., and LaLumiere, R.T. (2013). Posttraining optogenetic manipulations of basolateral amygdala activity modulate consolidation of inhibitory avoidance memory in rats. Proceedings of the National Academy of Sciences 110, 3597–3602. 10.1073/pnas.1219593110.

14. Blank, M., Dornelles, A.S., Werenicz, A., Velho, L.A., Pinto, D.F., Fedi, A.C., Schröder, N., and Roesler, R. (2014). Basolateral amygdala activity is required for enhancement of memory consolidation produced by histone deacetylase inhibition in the hippocampus. Neurobiol Learn Mem 111, 1–8. 10.1016/j.nlm.2014.02.009.

15. Lalumiere, R.T., Nguyen, L.T., and McGaugh, J.L. (2004). Post-training intrabasolateral amygdala infusions of dopamine modulate consolidation of inhibitory avoidance memory: involvement of noradrenergic and cholinergic systems. Eur J Neurosci 20, 2804–2810. 10.1111/j.1460-9568.2004.03744.x.

16. Kensinger, E.A., Garoff-Eaton, R.J., and Schacter, D.L. (2007). Effects of emotion on memory specificity: Memory trade-offs elicited by negative visually arousing stimuli. Journal of Memory and Language 56, 575–591. 10.1016/j.jml.2006.05.004.

17. Loftus, E.F., Loftus, G.R., and Messo, J. (1987). Some facts about “weapon focus.” Law Hum Behav 11, 55–62. 10.1007/BF01044839.

18. Wahlstrom, K.L., Huff, M.L., Emmons, E.B., Freeman, J.H., Narayanan, N.S., McIntyre, C.K., and LaLumiere, R.T. (2018). Basolateral Amygdala Inputs to the Medial Entorhinal Cortex Selectively Modulate the Consolidation of Spatial and Contextual Learning. J Neurosci 38, 2698–2712. 10.1523/JNEUROSCI.2848-17.2018.

19. Bass, D.I., Nizam, Z.G., Partain, K.N., Wang, A., and Manns, J.R. (2014). Amygdala-Mediated Enhancement of Memory for Specific Events Depends on the Hippocampus. Neurobiol Learn Mem 107, 37–41. 10.1016/j.nlm.2013.10.020.

20. Inman, C.S., Manns, J.R., Bijanki, K.R., Bass, D.I., Hamann, S., Drane, D.L., Fasano, R.E., Kovach, C.K., Gross, R.E., and Willie, J.T. (2018). Direct electrical stimulation of the amygdala enhances declarative memory in humans. Proc Natl Acad Sci USA 115, 98–103. 10.1073/pnas.1714058114.

21. Bass, D.I., and Manns, J.R. (2015). Memory-enhancing amygdala stimulation elicits gamma synchrony in the hippocampus. Behavioral Neuroscience 129, 244–256. 10.1037/bne0000052.

22. Manns, J.R., and Bass, D.I. (2016). The Amygdala and Prioritization of Declarative Memories. Curr Dir Psychol Sci 25, 261–265. 10.1177/0963721416654456.

23. Bass, D.I., Partain, K.N., and Manns, J.R. (2012). Event-specific enhancement of memory via brief electrical stimulation to the basolateral complex of the amygdala in rats. Behav Neurosci 126, 204–208. 10.1037/a0026462.

24. McGaugh, J.L. (2000). Memory--a Century of Consolidation. Science 287, 248–251. 10.1126/science.287.5451.248.

25. Roozendaal, B., Castello, N.A., Vedana, G., Barsegyan, A., and McGaugh, J.L. (2008). Noradrenergic activation of the basolateral amygdala modulates consolidation of object recognition memory. Neurobiol Learn Mem 90, 576–579. 10.1016/j.nlm.2008.06.010.

26. Hamann, S.B., Ely, T.D., Grafton, S.T., and Kilts, C.D. (1999). Amygdala activity related to enhanced memory for pleasant and aversive stimuli. Nat Neurosci 2, 289–293. 10.1038/6404.

27. McGaugh, J.L., and Roozendaal, B. (2009). Drug enhancement of memory consolidation: historical perspective and neurobiological implications. Psychopharmacology (Berl) 202, 3– 14. 10.1007/s00213-008-1285-6.

28. McGaugh, J.L. (2018). Emotional arousal regulation of memory consolidation. Current Opinion in Behavioral Sciences 19, 55–60. 10.1016/j.cobeha.2017.10.003.

29. Ahlgrim, N.S., and Manns, J.R. (2019). Optogenetic Stimulation of the Basolateral Amygdala Increased Theta-Modulated Gamma Oscillations in the Hippocampus. Front. Behav. Neurosci. 13, 87. 10.3389/fnbeh.2019.00087.

30. Tort, A.B.L., Komorowski, R.W., Manns, J.R., Kopell, N.J., and Eichenbaum, H. (2009). Theta–gamma coupling increases during the learning of item–context associations. PNAS 106, 20942–20947. 10.1073/pnas.0911331106.

31. Trimper, J.B., Stefanescu, R.A., and Manns, J.R. (2014). Recognition memory and theta– gamma interactions in the hippocampus. Hippocampus 24, 341–353. 10.1002/hipo.22228.

32. Wang, J., and Barbas, H. (2018). Specificity of Primate Amygdalar Pathways to Hippocampus. J. Neurosci. 38, 10019–10041. 10.1523/JNEUROSCI.1267-18.2018.

33. Pikkarainen, M., Rönkkö, S., Savander, V., Insausti, R., and Pitkänen, A. (1999). Projections from the lateral, basal, and accessory basal nuclei of the amygdala to the hippocampal formation in rat. J Comp Neurol 403, 229–260.

34. Vlcek, K., Fajnerova, I., Nekovarova, T., Hejtmanek, L., Janca, R., Jezdik, P., Kalina, A., Tomasek, M., Krsek, P., Hammer, J., et al. (2020). Mapping the Scene and Object Processing Networks by Intracranial EEG. Front. Hum. Neurosci. 14. 10.3389/fnhum.2020.561399.

35. Strange, B.A., Witter, M.P., Lein, E.S., and Moser, E.I. (2014). Functional organization of the hippocampal longitudinal axis. Nat Rev Neurosci 15, 655–669. 10.1038/nrn3785.

36. Witter, M.P., Wouterlood, F.G., Naber, P.A., and Van Haeften, T. (2000). Anatomical organization of the parahippocampal-hippocampal network. Ann N Y Acad Sci 911, 1–24. 10.1111/j.1749-6632.2000.tb06716.x.

37. Manns, J.R., and Eichenbaum, H. (2006). Evolution of declarative memory. Hippocampus 16, 795–808. 10.1002/hipo.20205.

38. Keller, C.J., Honey, C.J., Mégevand, P., Entz, L., Ulbert, I., and Mehta, A.D. (2014). Mapping human brain networks with cortico-cortical evoked potentials. Philos Trans R Soc Lond B Biol Sci 369, 20130528. 10.1098/rstb.2013.0528.

39. Friston, K.J. (1994). Functional and effective connectivity in neuroimaging: A synthesis. Human Brain Mapping 2, 56–78. 10.1002/hbm.460020107.

40. Friston, K.J. (2011). Functional and effective connectivity: a review. Brain Connect 1, 13–36. 10.1089/brain.2011.0008.

41. Büchel, C., Coull, J.T., and Friston, K.J. (1999). The predictive value of changes in effective connectivity for human learning. Science 283, 1538–1541. 10.1126/science.283.5407.1538.

42. Crocker, B., Ostrowski, L., Williams, Z.M., Dougherty, D.D., Eskandar, E.N., Widge, A.S., Chu, C.J., Cash, S.S., and Paulk, A.C. (2021). Local and distant responses to single pulse electrical stimulation reflect different forms of connectivity. Neuroimage 237, 118094. 10.1016/j.neuroimage.2021.118094.

43. Paulk, A.C., Zelmann, R., Crocker, B., Widge, A.S., Dougherty, D.D., Eskandar, E.N., Weisholtz, D.S., Richardson, R.M., Cosgrove, G.R., Williams, Z.M., et al. (2022). Local and distant cortical responses to single pulse intracranial stimulation in the human brain are differentially modulated by specific stimulation parameters. Brain Stimulation 15, 491–508. 10.1016/j.brs.2022.02.017.

44. Conner, C.R., Ellmore, T.M., DiSano, M.A., Pieters, T.A., Potter, A.W., and Tandon, N. (2011). Anatomic and electro-physiologic connectivity of the language system: a combined DTI-CCEP study. Comput Biol Med 41, 1100–1109. 10.1016/j.compbiomed.2011.07.008.

45. Matsui, T., Tamura, K., Koyano, K.W., Takeuchi, D., Adachi, Y., Osada, T., and Miyashita, Y. (2011). Direct comparison of spontaneous functional connectivity and effective connectivity measured by intracortical microstimulation: an fMRI study in macaque monkeys. Cereb Cortex 21, 2348–2356. 10.1093/cercor/bhr019.

46. Keller, C.J., Bickel, S., Entz, L., Ulbert, I., Milham, M.P., Kelly, C., and Mehta, A.D. (2011). Intrinsic functional architecture predicts electrically evoked responses in the human brain. Proceedings of the National Academy of Sciences 108, 10308–10313. 10.1073/pnas.1019750108.

47. Davis, T.S., Caston, R.M., Philip, B., Charlebois, C.M., Anderson, D.N., Weaver, K.E., Smith, E.H., and Rolston, J.D. (2021). LeGUI: A Fast and Accurate Graphical User Interface for Automated Detection and Anatomical Localization of Intracranial Electrodes. Front Neurosci 15, 769872. 10.3389/fnins.2021.769872.

48. Adamek, M., Swift, J.R., and Brunner, P. (2022). VERA -a versatile electrode localization framework. Version 1.0.0.

49. Bass, D.I., Partain, K.N., and Manns, J.R. (2012). Event-Specific Enhancement of Memory via Brief Electrical Stimulation to the Basolateral Complex of the Amygdala in Rats. Behav Neurosci 126, 204–208. 10.1037/a0026462.

50. Kundu, B., Davis, T.S., Philip, B., Smith, E.H., Arain, A., Peters, A., Newman, B., Butson, C.R., and Rolston, J.D. (2020). A systematic exploration of parameters affecting evoked intracranial potentials in patients with epilepsy. Brain Stimul 13, 1232–1244. 10.1016/j.brs.2020.06.002.

51. LaLumiere, R.T., McGaugh, J.L., and McIntyre, C.K. (2017). Emotional Modulation of Learning and Memory: Pharmacological Implications. Pharmacol Rev 69, 236–255. 10.1124/pr.116.013474.

52. McIntyre, C.K., Miyashita, T., Setlow, B., Marjon, K.D., Steward, O., Guzowski, J.F., and McGaugh, J.L. (2005). Memory-influencing intra-basolateral amygdala drug infusions modulate expression of Arc protein in the hippocampus. Proc Natl Acad Sci U S A 102, 10718–10723. 10.1073/pnas.0504436102.

53. Wahlstrom, K.L., Alvarez-Dieppa, A.C., McIntyre, C.K., and LaLumiere, R.T. (2021). The medial entorhinal cortex mediates basolateral amygdala effects on spatial memory and downstream activity-regulated cytoskeletal-associated protein expression. Neuropsychopharmacology 46, 1172–1182. 10.1038/s41386-020-00875-6.

54. Buchanan, T.W., and Adolphs, R. (2002). The role of the human amygdala in emotional modulation of long-term declarative memory. In Emotional cognition: From brain to behaviour Advances in Consciousness Research, vol. 44. (John Benjamins Publishing Company), pp. 9–34. 10.1075/aicr.44.02buc.

55. Duffley, G., Anderson, D.N., Vorwerk, J., Dorval, A.D., and Butson, C.R. (2019). Evaluation of methodologies for computing the deep brain stimulation volume of tissue activated. J Neural Eng 16, 066024. 10.1088/1741-2552/ab3c95.

56. Butson, C.R., Cooper, S.E., Henderson, J.M., and McIntyre, C.C. (2007). Patient-specific analysis of the volume of tissue activated during deep brain stimulation. NeuroImage 34, 661–670. 10.1016/j.neuroimage.2006.09.034.

57. Gunalan, K., Chaturvedi, A., Howell, B., Duchin, Y., Lempka, S.F., Patriat, R., Sapiro, G., Harel, N., and McIntyre, C.C. (2017). Creating and parameterizing patient-specific deep brain stimulation pathway-activation models using the hyperdirect pathway as an example. PLoS One 12, e0176132. 10.1371/journal.pone.0176132.

58. Howell, B., Choi, K.S., Gunalan, K., Rajendra, J., Mayberg, H.S., and McIntyre, C.C. (2018). Quantifying the axonal pathways directly stimulated in therapeutic subcallosal cingulate deep brain stimulation. Hum Brain Mapp 40, 889–903. 10.1002/hbm.24419.

59. Ríos, A.S., Oxenford, S., Neudorfer, C., Butenko, K., Li, N., Rajamani, N., Boutet, A., Elias, G.J.B., Germann, J., Loh, A., et al. (2022). Optimal deep brain stimulation sites and networks for stimulation of the fornix in Alzheimer’s disease. Nat Commun 13, 7707. 10.1038/s41467-022-34510-3.

60. Meyer, G.M., Hollunder, B., Li, N., Butenko, K., Dembek, T.A., Hart, L., Nombela, C., Mosley, P., Akram, H., Acevedo, N., et al. (2023). Deep Brain Stimulation for Obsessive-Compulsive Disorder: Optimal Stimulation Sites. Biological Psychiatry. 10.1016/j.biopsych.2023.12.010.

61. Horn, A., Reich, M.M., Ewert, S., Li, N., Al-Fatly, B., Lange, F., Roothans, J., Oxenford, S., Horn, I., Paschen, S., et al. (2022). Optimal deep brain stimulation sites and networks for cervical vs. generalized dystonia. Proc Natl Acad Sci U S A 119, e2114985119. 10.1073/pnas.2114985119.

62. Guillory, S.A., and Bujarski, K.A. (2014). Exploring emotions using invasive methods: review of 60 years of human intracranial electrophysiology. Soc Cogn Affect Neurosci 9, 1880–1889. 10.1093/scan/nsu002.

63. Parvizi, J., and Kastner, S. (2018). Promises and limitations of human intracranial electroencephalography. Nat Neurosci 21, 474–483. 10.1038/s41593-018-0108-2.

64. Richler, J.J., Palmeri, T.J., and Gauthier, I. (2013). How does using object names influence visual recognition memory? Journal of Memory and Language 68, 10–25. 10.1016/j.jml.2012.09.001.

65. Jun, S., Lee, S.A., Kim, J.S., Jeong, W., and Chung, C.K. (2020). Task-dependent effects of intracranial hippocampal stimulation on human memory and hippocampal theta power. Brain Stimulation 13, 603–613. 10.1016/j.brs.2020.01.013.

