## Supplemental Information for "Direct electrical stimulation of the human amygdala enhances recognition memory for objects but not scenes"

### Supporting Information

#### SI Material and Methods

##### Participants

Eight participants (62.5% Female, 37.5% Male; $M_{age} = 35.75$, $SD_{age} = 5.90$, $Range_{age} = 25\text{-}45$) with drug-resistant epilepsy volunteered to participate in the study. The study protocol was approved by the University of Utah and Washington University St. Louis Institutional Review Boards, and all participants provided written informed consent to all study procedures. To be included in the study subjects had to be English-speaking adults (> 18 y) implanted with intracranial depth electrodes, including those localized to the left or right amygdala. No exclusion was made based on gender, race, or ethnicity. Prior to intracranial monitoring, stereotactic EEG depth electrodes (Dixi/PMT: 0.80 mm diameter, 2.0 mm length platinum-coated contacts, spaced along 3.5 mm intervals; Ad-Tech: 1.3 mm diameter, 1.5 mm length platinum iridium-coated contacts, spaced along 5 mm intervals) were implanted into the brain parenchyma by a neurosurgeon for the sole purpose of clinical seizure investigation. All eight patients had electrode contacts localized near the BLA. The number of patients/sessions was predetermined by a formal power (target = 0.8) analysis that estimated the effect size of the main stimulation vs. no-stimulation memory effect based on effect sizes ranging from 0.88 to 1.02 (Cohen's *d* ) in three prior rat studies and one human study (G*Power)[1].

##### Brain Stimulation

*Prospective stimulation electrode localization.* Electrodes were localized using the LeGUI software package[2]. Briefly, preoperative MRIs were co-registered to post-implant CTs, and electrode contacts were detected via an automated density threshold. All automatic localizations were verified manually. LeGUI fits the localized electrodes to a standard space (Montreal Neurological Institute 152; MNI), from which the Neuro Morphometric atlas anatomical locations were derived. Atlas-derived amygdala locations were selected for stimulation.

*Memory testing stimulation parameters.* Stimulation parameters were chosen to replicate those used in three prior rat studies[3–5] and one human study[6] that showed amygdala-mediated memory enhancement. Specifically, stimulation was delivered to the BLA in bipolar, current-regulated, charge-balanced, biphasic rectangular pulses (500 us pulse width) at 1.0 mA (or 0.5 mA for three of the 14 sessions) for 1 s in eight trains of four pulses at 50 Hz. A research neurostimulator (CereStim M96; Blackrock Microsystems) combined with BCI2000[7] software were used to deliver stimulation precisely at the offset of image presentation for a randomized half of the encoding images. The current, duration, and pulse frequency are well below the typical clinical stimulation mapping parameters. A neurologist conducted a stimulation safety check prior to research testing and no epileptiform discharges were detected. Implanted electrodes were recorded continuously for clinical monitoring of seizure and epileptiform activity. No seizure activity or afterdischarges to stimulation were detected during research testing.

*Single-pulse stimulation parameters.* Single-pulse bipolar stimulation for the effective connectivity analyses was applied to the BLA as a single cathodic-first biphasic pulse (200 us pulse width) at 3 mA with a 1-3 s interpulse interval. A research neurostimulator (CereStim M96; Blackrock Microsystems) combined with either BCI2000[7] software or a custom software written in Labview (National Instruments, Austin, TX) were used to deliver single-pulse electrical stimulation. A return (or "ground") electrode was selected to minimize noise and stimulation artifact. One intracranial depth electrode in white matter was used to form a reference for recording. A Bovie grounding pad (McKesson, Irving, TX) was also placed on the patients' upper arm and attached to the research neurostimulator.

*Stimulation awareness test procedure.* Following the one-day retrieval test, a subset of participants were administered 10 stimulation and 10 sham stimulation (no current) trials delivered in random order to the BLA using a custom program in BCI2000[7]. Immediately after each trial participants were asked "Did you feel anything?" and were asked to use the left and right arrow keys on a keyboard to indicate "No" or "Yes", respectively. Both the experimenter and the participants were blind to which trials were paired with stimulation. Behavioral data were analyzed with GraphPad Prism 10. Stimulation detection accuracy (d') was calculated using estimates of the strength of the signal (endorsed "Yes" responses to stimulation trials; hits) relative to the noise (endorsed "Yes" responses to sham trials; false alarms)[8] (Fig. 3C).

**Stimuli**

During the encoding phase of the task, each participant viewed 180 images total (80 neutral object images presented on a scrambled background, 80 outdoor scene images, and 20 scrambled-only images). Outdoor scene images were taken from The Places dataset[9] and object images were taken from the Stark Lab's Mnemonic Similarity Task image-set[10,11]. All images were in color. Images were resized to ensure all images maintained the same number of pixels (400 x 400 pixels). The scrambled backgrounds were created using MATLAB (R2021b) function 'imscramble' with a maximum scrambling factor of 1 to scramble both object and scene images from the current study's image-set. Object images were centered onto a subset of these scrambled backgrounds and the remaining scrambled backgrounds were used as scrambled-only images (for controls in future electrophysiological analyses not within the scope of the present study). The Natural Image Statistical Toolbox[12] was used to ensure all images maintained a similar luminance, hue, and spatial frequency. To control for any potential salient features of the scrambled backgrounds themselves, patients were presented with object images on one version of a scrambled background during encoding and the same object image on a different scrambled background during retrieval.

**Analysis of Memory Performance**

Behavioral data were analyzed with GraphPad Prism 10. Recognition-memory performance was calculated using estimates of the strength of the signal (endorsed repeated/old images – i.e. hits) relative to the noise (endorsed new images – i.e. false alarms) (d')[6,8]. The d' was calculated for each experimental condition (stimulation vs. no stimulation) using MATLAB's dprime_simple function (norminv(hits)-norminv(false alarms)), with a correction for extreme proportions using the loglinear approach[13]. All data were normally distributed based on a Kolmogorov-Smirnov test of normality and no outliers were detected based on a Grubbs[14] test for each stimulation condition. One session resulted in a higher false alarm rate than hit rate for scene images resulting in negative d' values (Fig. 2A). In the present recognition-memory task patients were not only asked to mark images as 'old' or 'new', but also how certain they were in their response ('sure' or 'maybe'). Fig. S3 represents the analysis of these certainty judgments. Linear mixed effects modeling was performed for the one-day retrieval test for both the recognition memory accuracy (d') analyses (Fig. 2) and the memory confidence analyses (Fig. S3) to account for the multiple testing sessions within each patient (fixed effects for stimulation condition, random effects for patient and session, and memory performance as the dependent variable). To determine the effects of post-image presentation stimulation on the subsequent image, we also used linear mixed effects modeling to contrast the main amygdala-mediated memory-enhancement effect for object images with the memory performance for no-stimulation trials that occurred after a stimulation trial (Fig. S1). We also assessed the reaction times from image onset to participants' "old"/"new" key press response during the one-day retrieval test for all images. The average reaction time for a variety of image categories can be found in Fig. S2. Any image that produced a reaction time greater or less than two standard deviations from the mean was deemed an outlier and removed from the analyses. Reaction times were analyzed using either a *t* test or a one-way ANOVA with a Tukey's *post hoc* test. Fig. S4 displays the hit rates and false alarm rates from the recognition memory test for each participant/session. Hit and false alarm rates were analyzed using linear mixed effects modeling. Sessions were excluded from all analyses (including the electrophysiological analyses detailed below) if either the object or the scene false alarm rate exceeded 44%, a threshold based on expected false alarm rates from our previous work in a similar memory paradigm[6]. Sessions were also excluded if the delay between the encoding and retrieval session exceeded ~ 24 hours. (Before exclusion criteria: 17 patients total; 32 BLA stimulation sessions total.)

**Electrophysiological Data Analysis**
Neurophysiological data were recorded using a 128-channel data acquisition system (NeuroPort, Blackrock Microsystems, Salt Lake City, UT) or a Nihon Kohden JE-120 system (Nihon Kohden, Tokyo, Japan).

*Single-pulse stimulation analyses.* Analyses of electrophysiological data focused on evoked potentials recorded from the hippocampus in five patients for seven different BLA stimulation locations (the remaining three patients did not have the appropriate anterior and posterior hippocampal coverage and thus were not included in this analysis). The main question of interest was whether stimulation of the BLA reveals higher effective connectivity, as measured by the magnitude of evoked responses (SPEP), in the aHC important for object memory but not the pHC important for scene memory. Thus, each patient's hippocampus was parcellated into an anterior and posterior portion with separation at the uncal apex[15]. SPEP analyses were conducted with a custom Python script. Electrodes were bipolar re-referenced relative to nearest neighbors to account for volume conduction[16]. The data were filtered with a Butterworth bandpass filter (0.1 – 150 Hz). To attenuate 60 Hz electromagnetic noise in the evoked responses, 60 Hz and 120 Hz notch filters were applied. To analyze the SPEPs for the aHC and pHC data were divided into epochs spanning -900 to -10 ms prestimulation and 10-900 ms poststimulation (with stimulation onset at time zero) for each stimulation trial per hippocampal electrode. The number of trials ranged from 13 to 32 trials, median of 25 trials. We visually inspected the data and rejected trials with epileptiform spikes. The average number of trials removed across participants/sessions was 14.9 ± 12.8. For the remaining trials, voltages were normalized (Z-scored) to the prestimulation baseline data segment per stimulation trial. SPEP magnitude was measured as the normalized peak amplitude of the stimulation-evoked response in the 10-100 ms poststimulation period[17] using the trial-averaged waveform. Values in Fig. 4C represent the averaged evoked response across all anterior or all posterior hippocampal electrode contacts within a given patient/session. Linear mixed effects modeling was performed for the aHC and pHC comparison (Fig. 4C and 4D) to account for the multiple stimulation sessions within each patient (fixed effects for hippocampal location (aHC vs. pHC), random effects for patient and session, and SPEP amplitude as the dependent variable).

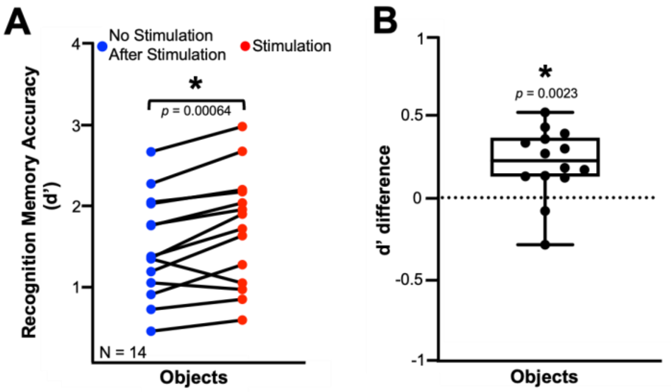


**Figure S1. Recognition-memory performance for objects followed by stimulation relative to no-stimulation objects that came after stimulation.** **A**, Recognition-memory performance (stimulation trials – no-stimulation trials after stimulation trials) for each session plotted as discriminability index (d') for objects. **B**, Recognition-memory performance plotted for each session as the difference in d' between the stimulation and no-stimulation-after-stimulation objects. If stimulation enhanced memory for subsequent no-stimulation images, then the data would show a decrease of the original memory-enhancement effect across sessions, but if stimulation diminished memory for subsequent no-stimulation images, then the data would show an increase of the original memory-enhancement effect across sessions. These results suggest no evidence of a change in our original effect and thus no significant carry-over effect of stimulation trials upon the subsequent encoding trials. *$p < 0.05$.

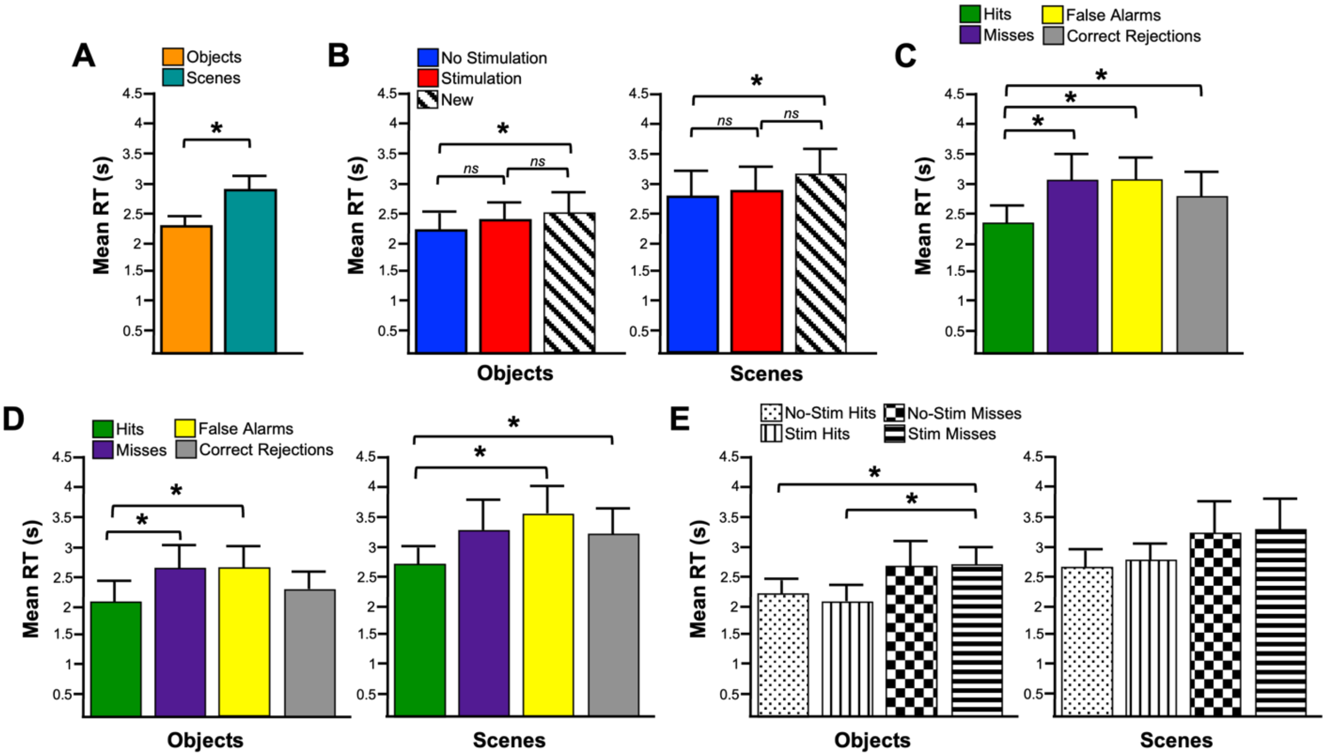


**Figure S2. Reaction times (RT) for recognition of previously viewed and new images at the one-day retrieval test. A**, Mean RTs across all retrieval sessions for previously viewed object and scene images. **B**, Mean RTs across all retrieval sessions for previously nonstimulated, previously stimulated, and new images for objects *(left)* and scenes *(right)*. **C**, Mean RTs across all retrieval sessions for Hits (correct "old" response for an old image), Misses (incorrect "new" response for an old image), False Alarms (incorrect "old" response for a new image), and Correct Rejections (correct "new" response for a new image). **D**, Mean RTs across all retrieval sessions for Hits, Misses, False Alarms, and Correct Rejections for objects *(left)* and scenes *(right)*. **E**, Mean RTs across all retrieval sessions for previously nonstimulated Hits, previously stimulated Hits, previously nonstimulated Misses, and previously stimulated Misses. *$p < 0.05$.

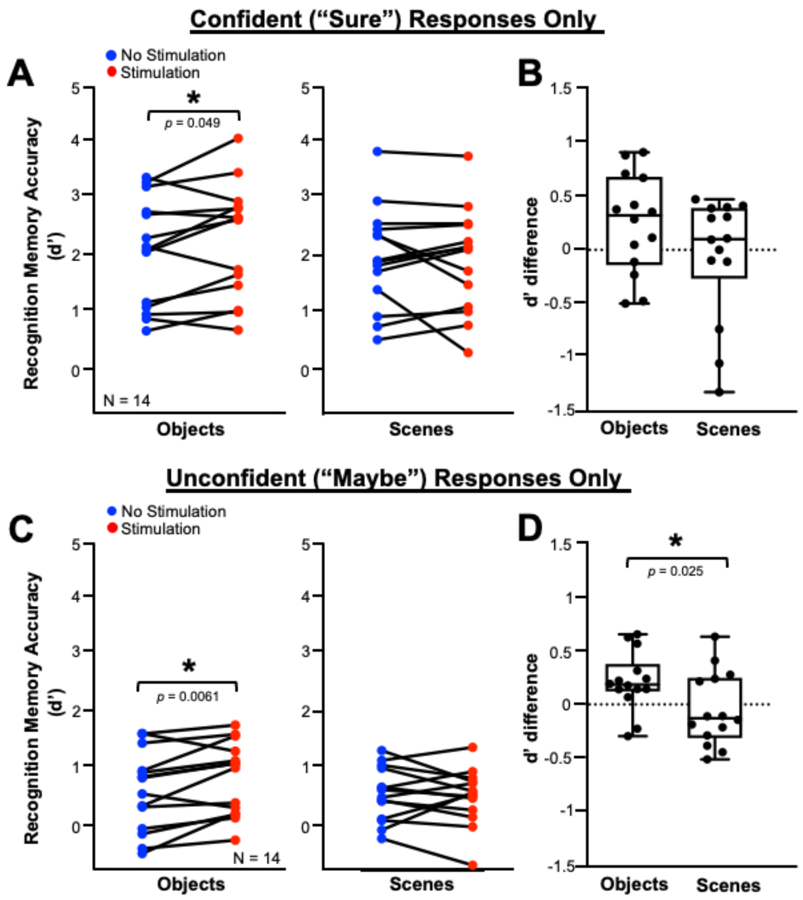


**Figure S3. BLA stimulation does not influence memory confidence. A**, Recognition-memory performance for each session plotted as discriminability index (d’) for object and scene images for “Sure” responses. **B**, Recognition-memory performance plotted for each session as the difference in d’ in the stimulation and no-stimulation conditions for the “Sure” responses (scatter plots in C). Overlaid box-and-whisker plots show the median, range, and interquartile range for each condition. **C**, Recognition-memory performance for each session plotted as discriminability index (d’) for object and scene images for “Maybe” responses. **D**, Recognition-memory performance plotted for each session as the difference in d’ in the stimulation and no-stimulation conditions for the “Maybe” responses (scatter plots in A). Overlaid box-and-whisker plots show the median, range, and interquartile range for each condition. *$p < 0.05$ compared with control images.

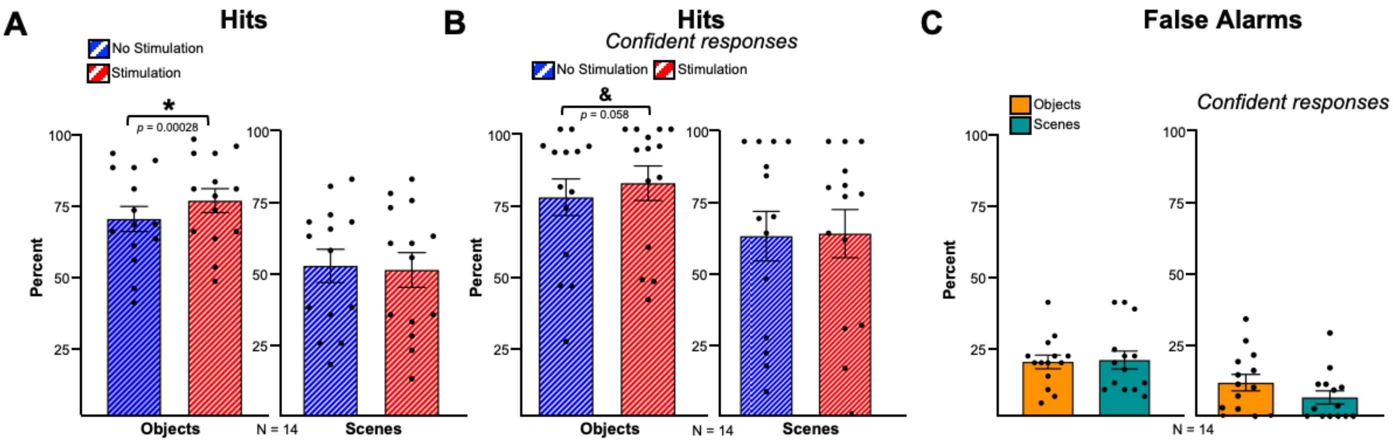


**Figure S4. Recognition memory performance for previously viewed images and new images during retrieval testing. A**, Number of responses in percent of Hits (correct "old" response for an old image) relative to the total number of old images for previously stimulated and nonstimulated object *(left)* and scene *(right)* images. **B**, Number of confident/"Sure" responses in percent of Hits relative to the total number of "Sure" responses for previously stimulated and nonstimulated object *(left)* and scene *(right)* images. This panel suggests the main results in panel A are essentially the same when we restrict the analyses to the putatively highest-quality data. **C**, Number of responses in percent of False Alarms (incorrect "old" response for a new image) relative to the total number of new images presented at retrieval testing *(left)*, and number of confident/"Sure" responses in percent of False Alarms relative to the total number of "Sure" responses *(right)*. *$p < 0.05$ ; &$p < 0.10$.

**Table S1. Participant demographics and relevant clinical neuropsychological testing scores**

| Subject | Sex | Age (yr) | Language dominance | FSIQ | VCI | PRI |
|---|---|---|---|---|---|---|
| 1 | Male | 32 | Left | 100 | 98 | 101 |
| 2 | Female | 38 | Unknown | Unknown | Unknown | Unknown |
| 3 | Female | 45 | Unknown | Unknown | Unknown | Unknown |
| 4 | Male | 40 | Unknown | 98 | 92 | 104 |
| 5 | Male | 37 | Unknown | 77 | Unknown | Unknown |
| 6 | Female | 25 | Left | 110 | 115 | 103 |
| 7 | Female | 34 | Left | 88 | 105 | 98 |
| 8 | Female | 35 | Left | 65 | 78 | 80 |

*All subjects reported being right hand-dominant. FSIQ, Full-Scale Intelligence Quotient; VCI, Verbal Comprehension Index; PRI, Perceptual Reasoning Index. Some information remains inaccessible from patient medical records and is therefore listed as "Unknown".*

**Table S2. General epilepsy patient information**

| Subject | Seizure focus | Preoperative image findings | AEDs during testing |
|---|---|---|---|
| 1 | R hippocampus/amygdala/entorhinal cortex and L hippocampus | Potential small cyst in the R medial temporal lobe | Lorazepam, Brivaracetam, Levetiracetam |
| 2 | R temporal pole/gyrus, L temporal gyrus, B/l hippocampus | Hypometabolism involving B/l medial and anterior temporal lobes | Eslicarbazepine, Clonazepam |
| 3 | B/l hippocampus | Encephalocele arising from the medial R temporal lobe | Zonisamide, Lorazepam, Cenobamate |
| 4 | L hippocampus/entorhinal cortex | L hippocampus is slightly under rotated | Lamotrigine |
| 5 | R hippocampus/amygdala | L temporal cortical malformation involving the hippocampus and amygdala | Levetiracetam, Lamotrigine, Clobazam, Lorazepam |
| 6 | L temporal and occipital gyri | L temporo-occipital junction white matter cortical dysplasia | Zonisamide, Levetiracetam, Lacosamide, Clobazam, Cenobamate |
| 7 | R amygdala/hippocampus/orbitofrontal/middle temporal gyrus | Normal | Clobazam, Lorazepam, Lamotrigine, Levetiracetam |
| 8 | R mesial temporal lobe (hippocampus) and lateral temporal lobe | Slightly decreased volume of the R temporal lobe | Oxcarbazepine |

*AEDs, anti-epileptic drugs; B/l, bilateral; L, left; R, right.*

**Table S3. BLA Stimulation Location**

| Subject | Stimulation session | Stimulation hemisphere | Medial or lateral BLA |
|---|---|---|---|
| 1 | 1 | Right | Medial |
| | 2 | Right | Lateral |
| | 3 | Right | Medial |
| 2 | 1 | Right | Lateral |
| | 2 | Left | Lateral |
| 3 | 2 | Left | Lateral |
| 4 | 1 | Left | Medial |
| | 2 | Right | Lateral |
| 5 | 1 | Left | Lateral |
| 6 | 1 | Left | Lateral |
| | 2 | Left | Medial |
| 7 | 1 | Right | Medial |
| | 2 | Right | Lateral |
| 8 | 1 | Right | Medial |

*Amygdala location stimulated for each patient and stimulation session within patient.*

**Table S4. Stimulation d' and d' differences at the one-day retrieval test**

| Subject | Session | Objects no stimulation | Objects stimulation | Objects difference | Scenes no stimulation | Scenes stimulation | Scenes difference |
|---|---|---|---|---|---|---|---|
| 1 | 1 | 0.938 | 0.877 | -0.061 | 0.412 | 0.572 | 0.16 |
| | 2 | 0.569 | 0.755 | 0.186 | 0.646 | 0.227 | -0.419 |
| | 3 | 0.495 | 0.495 | 0 | 0.796 | 0.732 | -0.064 |
| 2 | 1 | 1.6 | 1.96 | 0.358 | 0.646 | 0.917 | 0.271 |
| | 2 | 1.96 | 2.1 | 0.142 | 0.695 | 0.495 | -0.2 |
| 3 | 2 | 2.19 | 2.61 | 0.422 | 0.828 | 0.902 | 0.074 |
| 4 | 1 | 1.31 | 1.64 | 0.323 | 2.27 | 1.75 | -0.529 |
| | 2 | 0.828 | 0.957 | 0.128 | 1.48 | 1.29 | -0.191 |
| 5 | 1 | 1.62 | 1.88 | 0.263 | 1.24 | 1.47 | 0.223 |
| 6 | 1 | 1.44 | 1.82 | 0.38 | 0.902 | 1.47 | 0.568 |
| | 2 | 2.48 | 2.92 | 0.44 | 1.73 | 1.73 | 0 |
| 7 | 1 | 1.55 | 1.55 | 0 | 0.657 | 0.594 | -0.063 |
| | 2 | 0.693 | 1.19 | 0.497 | -0.0626 | -0.326 | -0.2634 |
| 8 | 1 | 1.99 | 2.13 | 0.139 | 1.26 | 1.13 | -0.131 |

*All measures use d prime (d') which is calculated as the normalized hit rate minus false alarm rate.*

1. Erdfelder, E., Faul, F., and Buchner, A. (1996). GPOWER: A general power analysis program. Behavior Research Methods, Instruments, & Computers *28*, 1–11. https://doi.org/10.3758/BF03203630.

2. Davis, T.S., Caston, R.M., Philip, B., Charlebois, C.M., Anderson, D.N., Weaver, K.E., Smith, E.H., and Rolston, J.D. (2021). LeGUI: A Fast and Accurate Graphical User Interface for Automated Detection and Anatomical Localization of Intracranial Electrodes. Front Neurosci *15*, 769872. https://doi.org/10.3389/fnins.2021.769872.

3. Bass, D.I., and Manns, J.R. (2015). Memory-enhancing amygdala stimulation elicits gamma synchrony in the hippocampus. Behavioral Neuroscience *129*, 244–256. https://doi.org/10.1037/bne0000052.

4. Bass, D.I., Nizam, Z.G., Partain, K.N., Wang, A., and Manns, J.R. (2014). Amygdala-Mediated Enhancement of Memory for Specific Events Depends on the Hippocampus. Neurobiol Learn Mem *107*, 37–41. https://doi.org/10.1016/j.nlm.2013.10.020.

5. Bass, D.I., Partain, K.N., and Manns, J.R. (2012). Event-Specific Enhancement of Memory via Brief Electrical Stimulation to the Basolateral Complex of the Amygdala in Rats. Behav Neurosci *126*, 204–208. https://doi.org/10.1037/a0026462.

6. Inman, C.S., Manns, J.R., Bijanki, K.R., Bass, D.I., Hamann, S., Drane, D.L., Fasano, R.E., Kovach, C.K., Gross, R.E., and Willie, J.T. (2018). Direct electrical stimulation of the amygdala enhances declarative memory in humans. Proc Natl Acad Sci USA *115*, 98–103. https://doi.org/10.1073/pnas.1714058114.

7. Schalk, G., McFarland, D.J., Hinterberger, T., Birbaumer, N., and Wolpaw, J.R. (2004). BCI2000: a general-purpose brain-computer interface (BCI) system. IEEE Trans Biomed Eng *51*, 1034–1043. https://doi.org/10.1109/TBME.2004.827072.

8. Macmillan, N.A., and Kaplan, H.L. (1985). Detection theory analysis of group data: estimating sensitivity from average hit and false-alarm rates. Psychol Bull *98*, 185–199.

9. Zhou, B., Lapedriza, A., Khosla, A., Oliva, A., and Torralba, A. (2018). Places: A 10 Million Image Database for Scene Recognition. IEEE Trans Pattern Anal Mach Intell *40*, 1452–1464. https://doi.org/10.1109/TPAMI.2017.2723009.

10. Stark, S.M., Stevenson, R., Wu, C., Rutledge, S., and Stark, C.E.L. (2015). STABILITY OF AGE-RELATED DEFICITS IN THE MNEMONIC SIMILARITY TASK ACROSS TASK VARIATIONS. Behavioral neuroscience *129*, 257. https://doi.org/10.1037/bne0000055.

11. Yassa, M.A., Lacy, J.W., Stark, S.M., Albert, M.S., Gallagher, M., and Stark, C.E. (2010). Pattern separation deficits associated with increased hippocampal CA3 and dentate gyrus activity in nondemented older adults. Hippocampus *21*, 968. https://doi.org/10.1002/hipo.20808.

12. Bainbridge, W.A., and Oliva, A. (2015). A toolbox and sample object perception data for equalization of natural images. Data in Brief *5*, 846–851. https://doi.org/10.1016/j.dib.2015.10.030.

13. Hautus, M.J. (1995). Corrections for extreme proportions and their biasing effects on estimated values ofd′. Behavior Research Methods, Instruments, & Computers *27*, 46–51. https://doi.org/10.3758/BF03203619.

14. Grubbs, F.E. (1950). Sample criteria for testing outlying observations. Annals of Mathematical Statistics *21*, 27–58. https://doi.org/10.1214/aoms/1177729885.

15. Inman, C.S., James, G.A., Vytal, K., and Hamann, S. (2018). Dynamic changes in large-scale functional network organization during autobiographical memory retrieval. Neuropsychologia *110*, 208–224. https://doi.org/10.1016/j.neuropsychologia.2017.09.020.

16. Paulk, A.C., Zelmann, R., Crocker, B., Widge, A.S., Dougherty, D.D., Eskandar, E.N., Weisholtz, D.S., Richardson, R.M., Cosgrove, G.R., Williams, Z.M., et al. (2022). Local and distant cortical responses to single pulse intracranial stimulation in the human brain are differentially modulated by specific stimulation parameters. Brain Stimulation *15*, 491–508. https://doi.org/10.1016/j.brs.2022.02.017.

17. Kundu, B., Davis, T.S., Philip, B., Smith, E.H., Arain, A., Peters, A., Newman, B., Butson, C.R., and Rolston, J.D. (2020). A systematic exploration of parameters affecting evoked intracranial potentials in patients with epilepsy. Brain Stimul *13*, 1232–1244. https://doi.org/10.1016/j.brs.2020.06.002.
